# Novel Endogenous Retrovirus in the Slow Loris

**DOI:** 10.1101/2025.01.02.631053

**Authors:** Charles Michie, Hayley Beth Free, Vincent Nijman, Ravinder Kanda

## Abstract

Endogenous retroviruses (ERVs) are the result of an exogenous infectious retrovirus becoming integrated within the host genome through infection of germline cells. The majority of ERV research has been conducted on humans and other great apes and research of them within other primates can provide unique insights. Screening the genome of two endangered slow lorises *Nycticebus bengalensis* and *N. coucang* with a previously identified LTR, a novel ERV was identified within their genomes with varying levels of completeness and numerous solo LTRs. Phylogenetic analysis of the genes of this ERV indicates that it is a betaretrovirus most closely related to HERV-K. Despite being closely related to HERV-K, the ERV does not appear to be within humans and is only found in Asian lorises and absent in all other primates. Due to the staggering similarity of viral insertions of this ERV between the two species of slow lorises, indicates that the current *N. coucang* genome may actually be a hybrid of the two species or a *N. bengalensis* from a different population as the reference genome. This study highlights that studying ERVs in more distant primates provides information about the genomes and evolution of these species as well as viral evolution.

## 1 Introduction

Endogenous retroviruses (ERVs) are genomic remnants of ancient exogenous retroviral infections, that became endogenized into vertebrate genomes through infection of germline cells, allowing their inheritance in a mendelian fashion (Weiss, 2006). While most ERVs do not become fixed in a species population, those that do can constitute a significant proportion of the host genome, where in humans approximately 8% of our genome are ERVs (Lander et al., 2001). ERVs were long considered to be “junk DNA”, non-functional elements occupying space in the genome, but further study has demonstrated that they can have large impacts; contributing to novel genes, acting as promoters, disrupting genes and having links to various diseases (Chen et al., 2024; Katzourakis et al., 2005; Xue et al., 2020). The structure of ERVs typically consists of two long terminal repeats (LTRs) with four major genes between: gag, pro, pol and *env* (Coffin, 1992). By studying these ERVs we can gain information about the retroviruses which infected past populations, and the impact of these insertions on the genome evolution of the host species.

The study of ERVs in primates genomes has mainly focused on humans and the other great apes. There has been a particular focus on human endogenous retroviruses (HERVs) and although all HERV families have been identified in chimp genomes, some HERV loci are unique to humans, suggesting integration of these particular insertions relatively recently (Li et al., 2022). The study of ERVs within humans and great apes genomes has been beneficial in investigating host-virus interactions, as well as how ERVs have contributed to their genomics and disease (Grandi et al., 2019; Grandi & Tramontano, 2017). Comparatively very little study of ERVs has been accomplished in the strepsirrhines, with what studies that have been undertaken within strepsirrhines largely focused on one superfamily, the lemurs (Lemuroidea) with very few little on lorises and bushbabies (Lorisoidea).

The Lorisoidea are small nocturnal primates found in Africa (Galagidae) and Asia (Lorisidae). The Asian lorises comprise three genera, i.e., slow lorises (*Nycticebus*), pygmy lorises (*Xanthonycticebus*) and slender lorises (*Loris*) (Nekaris, 2014; Nekaris & Nijman, 2022; Nekaris & Starr, 2015). All lorises species are on the IUCN Red List (status varying from Vulnerable to Critically Endangered), largely due to deforestation and trade for pets or medicinal purposes (Moore et al., 2014; Nekaris & Burrows, 2020). To date there has been very little study of ERVs within lorises, with only two studies describing ERVs within the slow loris (Gifford et al., 2005; Li et al., 2022). Studies of retroviruses that infected and evolved with lorises may provide an opportunity to understand the basal traits of primate retroviruses. By screening within smaller primates, we can examine if patterns seen in larger primates which have been extensively studied such as number of loci, families are the same in smaller primates and possibly if there are unique adaptations within lorises due to retroviral insertions.

Here, we use an LTR (Nycticebus_bengalensis_LG02 100652008 – 100652802 Unknown_HERV_3LTR) identified in Li et al., (2022) from an “Unknown HERV” to scan for new viruses within two species, the Bengal slow loris *Nycticebus bengalensis* and the greater slow loris *N. coucang*. Although they report a single retroviral insertion it did not have all four major genes. We are interested in understanding the evolutionary dynamics of ERVs in *Nycticebus* genomes. By comparing viral insertions in these closely related species it is possible to study how the ERV has changed within the genome since species split and if there have been any genomic events.

## 2 Methods

### 2.1 Genome screening

To investigate the number and distribution of loci of the novel ERV identified by Li et al., (2022), the 715 bp sequence “Nycticebus_bengalensis_LG02 100652008 – 100652802 Unknown_HERV_3LTR” (Supplementary Figure 1) was extracted from the *N. bengalensis* reference genome and used as a query in BLASTn (default parameters) to perform in silico screening of the reference genomes of *Nycticebus bengalensis* (GCA_023898255.1) and *N. coucang* (GCA_027406575.1).

The LTR BLASTn hits were used to identify complete and incomplete full-length ERVs, truncated viral elements, and solo LTRs. Any BLASTn hit that was over 400 bp long and had an e-value of less than 1×10^-5^ was considered a real LTR and analysed further. The LTR BLASTn hits which were longer than 400 bp but were incomplete (<715 bp), we calculated the expected location of the start and end of the LTR based on difference between query and subject length in the genome and extracted these.

### 2.2 Identification of Full-length ERVs

Potential full-length ERVs were determined by identifying LTRs that were within 20kb bases of each other, on the same chromosome, and in the same orientation. These pairs of LTRs were extracted and aligned manually in Geneious Prime v2024.0.3 (https://www.geneious.com/). To identify the internal genes of this ERV, we focused on those pairs of LTRs that had high sequence similarity (>94%) and were therefore more likely to represent recent insertions with intact internal genes (LTRs of a full-length insertion are identical at the time of integration and become more divergent with time as they accumulate mutations). For the pairs of LTRs that satisfied these criteria, the internal region between the LTRs was extracted, and open reading frames were identified using the NCBI ORFfinder (https://www.ncbi.nlm.nih.gov/orffinder/). A BLASTp search against the non-redundant protein sequences database (restricted to Viruses – taxid: 10239) was used to identify the gag, pro, pol, and env genes. Sequences which had all four core genes (gag, pro, pol, env) and did not have large insertions (over 2000 bases) were aligned in Geneious (MUSCLE v5.1; Edgar, 2004). As none of the insertions identified had the four complete internal genes, we created a consensus sequence of each gene from the full length elements we identified, to enable comparison to other ERVs. A number of insertions were missing one or more of the core genes – these we term incomplete full-length ERVs. Conserved domains were identified with the conserved domain database (Wang et al., 2023).

### 2.3 Phylogenetic analysis of the full-length ERV

The phylogenetic history of the constructed consensus full-length ERV was determined by alignment of conserved domains within each gene to increase strength of alignment. Separate phylogenetic trees were made for each gene. Those used were the p24 N region of *gag*, RVP of *pro*, RT of *pol* and TM of *env* translated amino acids with representative retroviruses from several retroviral families (Supplementary Table 1). The retroviruses compared against were biassed towards Betaretroviruses as the best BLAST hits were Betaretroviruses. Epsilonretroviruses were excluded from all analyses, and Spumaretroviruses were excluded in *env* analysis as they resulted in poor alignments. Only Alpharetroviruses, Betaretroviruses and Deltaretroviruses have a p24 conserved domain in *gag* therefore only these were included in the *gag* phylogeny. The position of the p24, RVP and RT was determined by the conserved domain database (Wang et al., 2023), whereas the TM was determined by the start of the cleavage site (R-X-R/K-R) until the end of the *env* (Bénit et al., 2001). All regions were aligned in Geneious (MUSCLE). ModelFinder (Kalyaanamoorthy et al., 2017) on IQTree v2.3.5 (Minh et al., 2020) was used to determine the best model of protein evolution (*gag* – LG+I+G4, *pro* and *pol* – rtREV+G4, *env* – LG+G4).

Maximum likelihood trees were constructed with PhyML v3.3. 20180621 (Guindon et al., 2010) with 1000 bootstraps and Bayesian trees were constructed with MrBayes v3.26 (Huelsenbeck & Ronquist, 2001) with a burn-in length of 100,000 and 1,000,000 Markov chain Monte Carlo (MCMC) steps with four heated chains and heated chain temperature of 0.2 with a sampling frequencing of 200 trees. Final effective sample sizes of 1812, 1435, 1319, 1440 for *gag, pro pol* and *env* analyses respectively indicated convergence of the Bayesian MCMC analysis. Trees were visualised with TreeViewer v2.2 (Bianchini & Sánchez-Baracaldo, 2024).

### 2.4 LTR insertion categorisation

For the remaining LTRs, an additional 1kb of flanking sequence from the 5’ and 3’ end of each insertion was extracted to identify whether those loci represent true solo LTRs, or 5’/3’ truncated FL elements (Figure 1). The consensus gag and env gene were used to BLASTn these extended LTRs. In rare cases there were triplets, where an LTR was associated with two viral elements.

**Figure 1.**
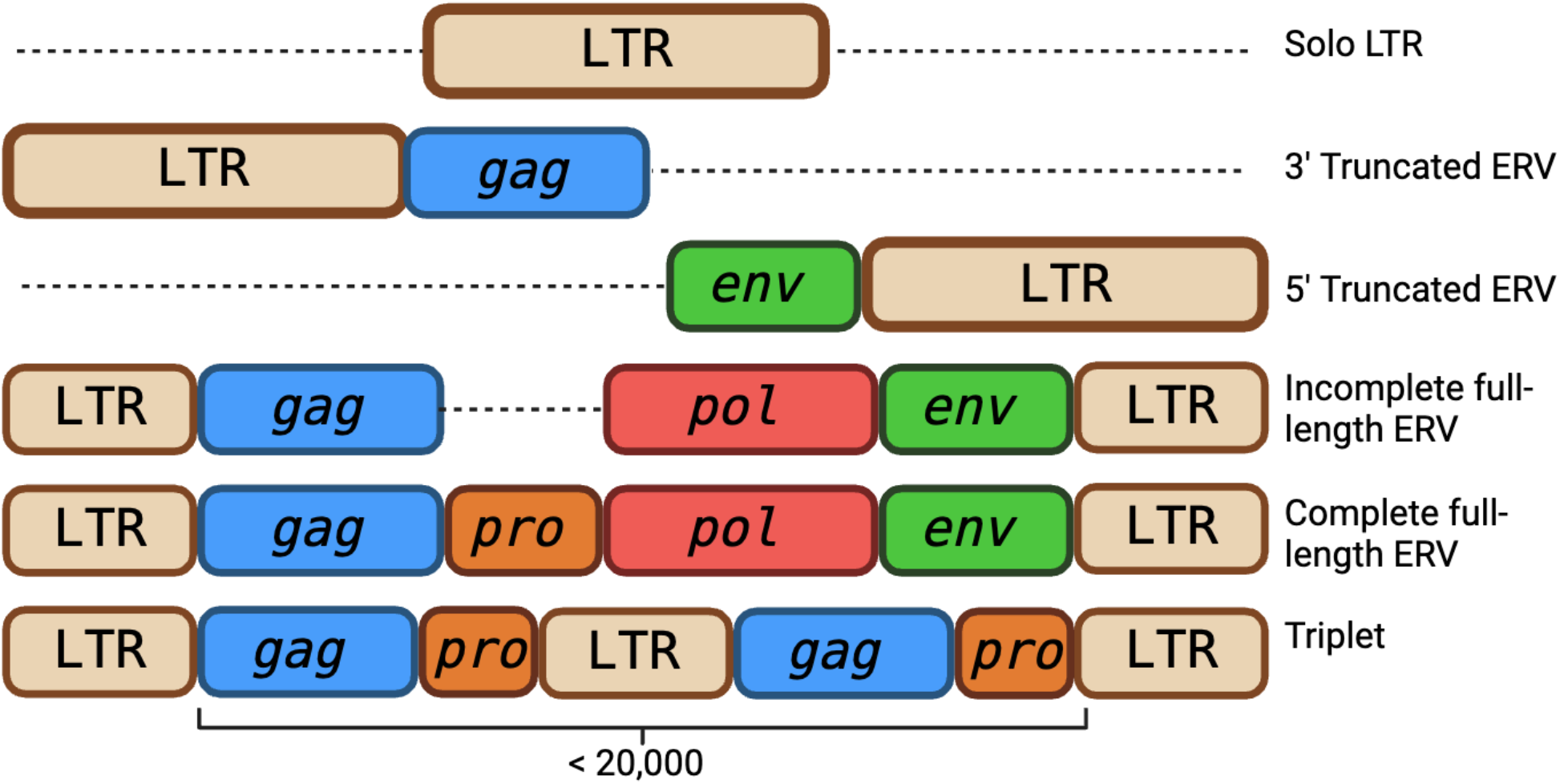
Graphical representation of categories of novel retroviral insertions identified within the genomes of *Nycticebus coucang* and *Nycticebus bengalensis*.

### 2.5 Similarity of the novel full-length ERV in *Nycticebus coucang* reference genome

The full length ERV loci identified in either *N. bengalensis* or *N. coucang* were compared to determine if these insertions predated the divergence of these species (i.e. present in the same state in both species), or detect insertional polymorphism (i.e. FL in one species, but solo LTR/preinsertion site in the other species) which would indicate relatively recent activity of this ERV (i.e after these two species had diverged). To determine if the viral insertion was the same between the two species, the target site duplications (TSDs) and flanking region of each full-length pair and truncated LTR found within the *N. bengalensis* was compared against those found in the *N. coucang*. If the TSDs between species were identical and the insertion was on the same chromosome it was determined to be the same viral insertion. The order of chromosomes was different between the reference genomes, therefore segments of each chromosome from each genome was compared against the opposing genome using BLASTn to match the chromosomes together (Supplementary Table 2). As there was no Y chromosome available for *N. bengalensis* this was excluded.

The similarity between all insertions (excluding solo LTRs) was determined through alignment in Geneious (MUSCLE) and the percentage of sites that were identical was found. The similarity was determined between the whole insertion and between the 5’ and 3’ LTRs in the genomes of the two species. The number of pairwise differences was also calculated.

### 2.6 Estimation of age of insertions

The age of insertion was estimated from the amount of divergence between LTRs. This was only calculated for full length ERVs in *N. coucang* as the *N. bengalensis* genome has regions of lower quality. Time was calculated as T = D/(μ * L), where D is the number of differences between the two LTRs, μ is the mutation rate, and L is the length of the LTR. 1.72×10^-9^ (per bp per year) is used as the mutation rate from Lemurs (Campbell et al., 2021). Clear large indels were ignored and removed from the alignment when calculating differences between the LTRs as they cannot be accounted for by the neutral evolutionary mutation rate.

### 2.7 Identification of the novel full-length ERV in other primates

To identify what other primate genomes the full-length ERV is present in, the consensus *env* gene and 3’ LTR (2672bp) were screened with BLASTn against the reference genomes of eight other strepsirrhine with varying amounts of divergence from *N. bengalensis* as representatives of other clades and the reference human genome (*Xanthonycticebus pygmaeus* - GWHBCHX00000000; *Loris tardigradus* - GCA_023783135; *L. lydekkerianus* - GCA_963574355; *Perodicticus potto* - GCA_963574655; *Otolemur garnettii -* GCA_000181295; *Lemur catta* - GCA_020740605; *Homo sapiens -* GCF_000001405). If there were any BLAST hits which had higher than 80% percent identity and 75% query coverage (2004bp) then it was determined that the ERV was also present in that species.

## 3 Results

### 3.1 Identification of a full-length ERV in *Nycticebus bengalensis*

Screening the *N. bengalensis* genome with the LTR from (Li et al., 2022) there were 1498 hits over 400bp long, of which 184 were within 20kp of each other creating 96 pairs as some LTRs were involved into multiple pairs. After removal of pairs which contained an LTR that did not have a clear start or end motif or contained indels there were 84 pairs of LTRs in total (Supplementary Table 3). Of these 84 pairs, 27 had a pairwise identity above 94%. After finding open reading frames and BLASTp of these, there were 16 sequences which contained all four genes, 14 of which did not have large insertions and were used to construct a consensus full-length ERV which is 8923bp long (NCBI Accession numbers to be provided). This consensus full-length ERV was mostly complete, where only six indels or bases changes were needed to create full length genes. Each gene was in a different reading frame.

#### 3.1.1 Conserved domains in the full-length ERV

The full-length ERV contained many of the typical conserved domains for retroviruses. The *gag* gene contained a myristylation signal at the beginning, followed by a truncated p10*gag*, an N and C terminal p24*gag*, and a single zinc finger. In the *pro* gene there was a dUTPase at the beginning typical of betaretroviruses (Hizi & Herzig, 2015), which is followed the protease which contains the catalytic motif of DSG which is more typical of alpharetroviruses, whereas DTG is common in betaretroviruses (Chameettachal et al., 2023; Konvalinka et al., 2015). The *pol* begins with the reverse transcriptase (RT) which has a catalytic site of YMDD. There is only one heptad repeat in the *env*.

### 3.2 Phylogenetic analysis

Both bayesian and maximum likelihood analysis of all four major genes indicated that the novel ERV is a betaretroviruses, most closely related to HERV-K (Figure 2; Supplementary Figure 2). Following similar nomenclature, this virus has provisionally been named Loris endogenous retrovirus 1 (LERV1). Notably, LERV1 is distinct from the one other ERV identified in the *Nycticebus* genus, RV Slow Loris (Gifford et al., 2005).

**Figure 2.**
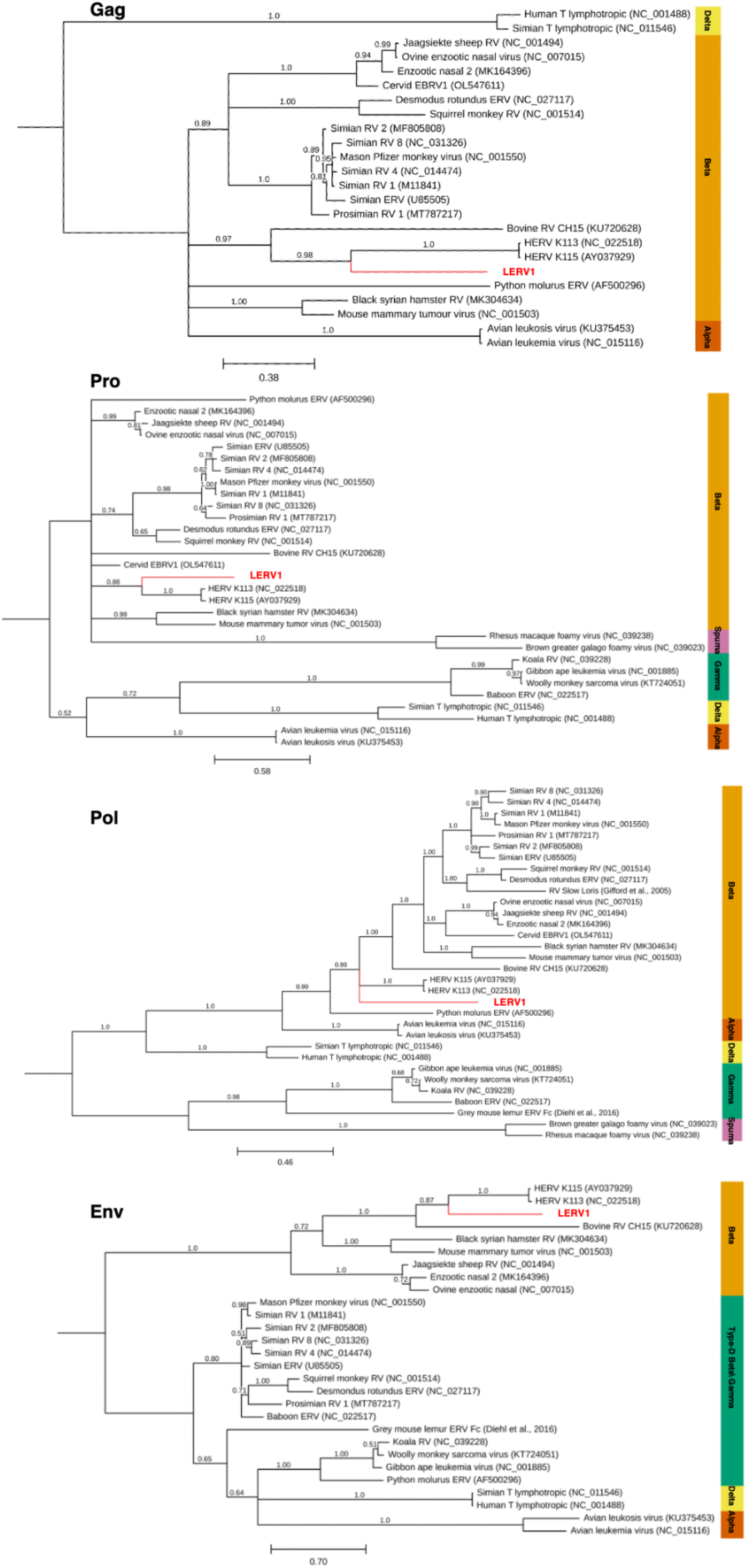
Phylogeny of LERV1 (in red) inferred from Bayesian (MrBayes) analysis with posterior probability shown. Scale bar represents substitutions per site. Trees have been rooted at midpoint. Maximum likelihood inference (PhyML) can be found in Supplementary Figure 2.

### 3.3 LTR identification and categorization in the two *Nycticebus* genomes

In *N. coucang* there were a total of 1426 solo LTRs and 45 complete full length elements, whilst in *N. bengalensis* there were 1305 solo LTRs and 33 complete full length elements. In both species the most viral elements were on Chromosomes 1 and X (Figure 3). No significant difference was found between the number of insertions and the distribution across the genome between species (ANOVA: p-value = 0.71). There was no bias towards orientation in either species (Chi-Squared Test: *N. bengalensis* p-value = 0.17, *N. coucang* p-value = 0.66). An insertion with three LTRs, referred to as triplets, represent full-length insertions where there has been a duplication of an LTR and internal genes resulting in an insertion with three LTRs. These were more common within *N. bengalensis* where there has been multiple duplications of the entire insertion whilst in *N. coucang* there is a single copy of the insertion (Figure 4A). The disparity between number of LTR hits and number of LTRs involved in pairs can be partly explained by difference in chromosome length and likely being missed in assembly (Supplementary Table 2), whilst others can be explained by a genomic event within one genome (Supplementary Table 3). For example, within the *N. bengalensis* a complete full-length pair (*N. coucang* X:124248563-124259123) has undergone homologous recombination and is now a solo LTR (*N. bengalensis* X:122693788-122696469) as evident by identical TSDs and flanking regions (Figure 4B). In both species there is evidence of inversions on chromosomes where a full-length element is in different orientations in the two genomes. 10 full-length insertions were unique to the *N. coucang* genome; however, we cannot be confident that this represents LERV1 activity after the divergence of these two species, as the pre-integration site was absent from the *N. bengalensis* genome implying that the whole region may be missing from the assembly.

**Figure 3.**
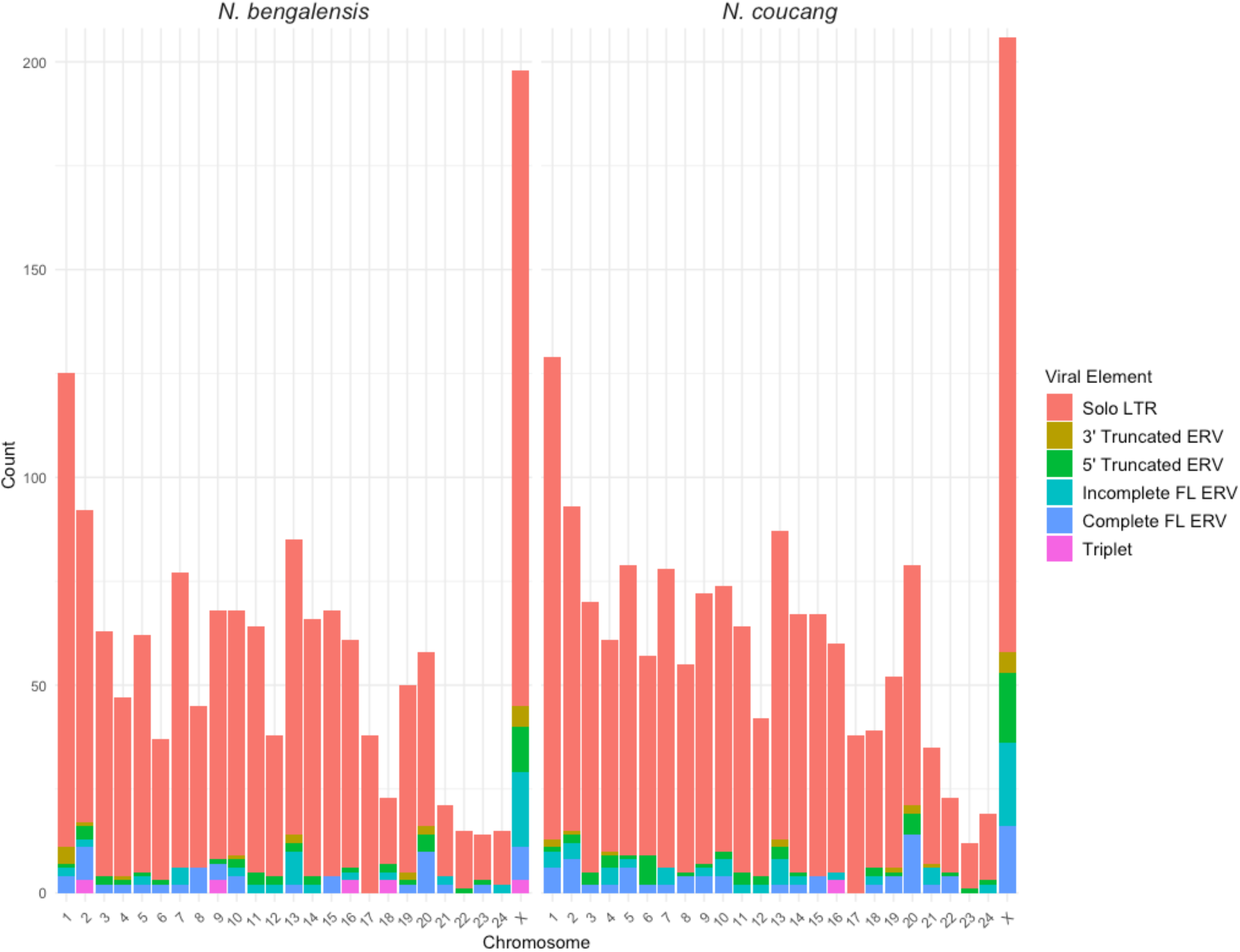
Number of LERV1 LTR insertions according to insertion category in *N. bengalensis* and *N. coucang* reference genomes across every chromosome.

**Figure 4.**
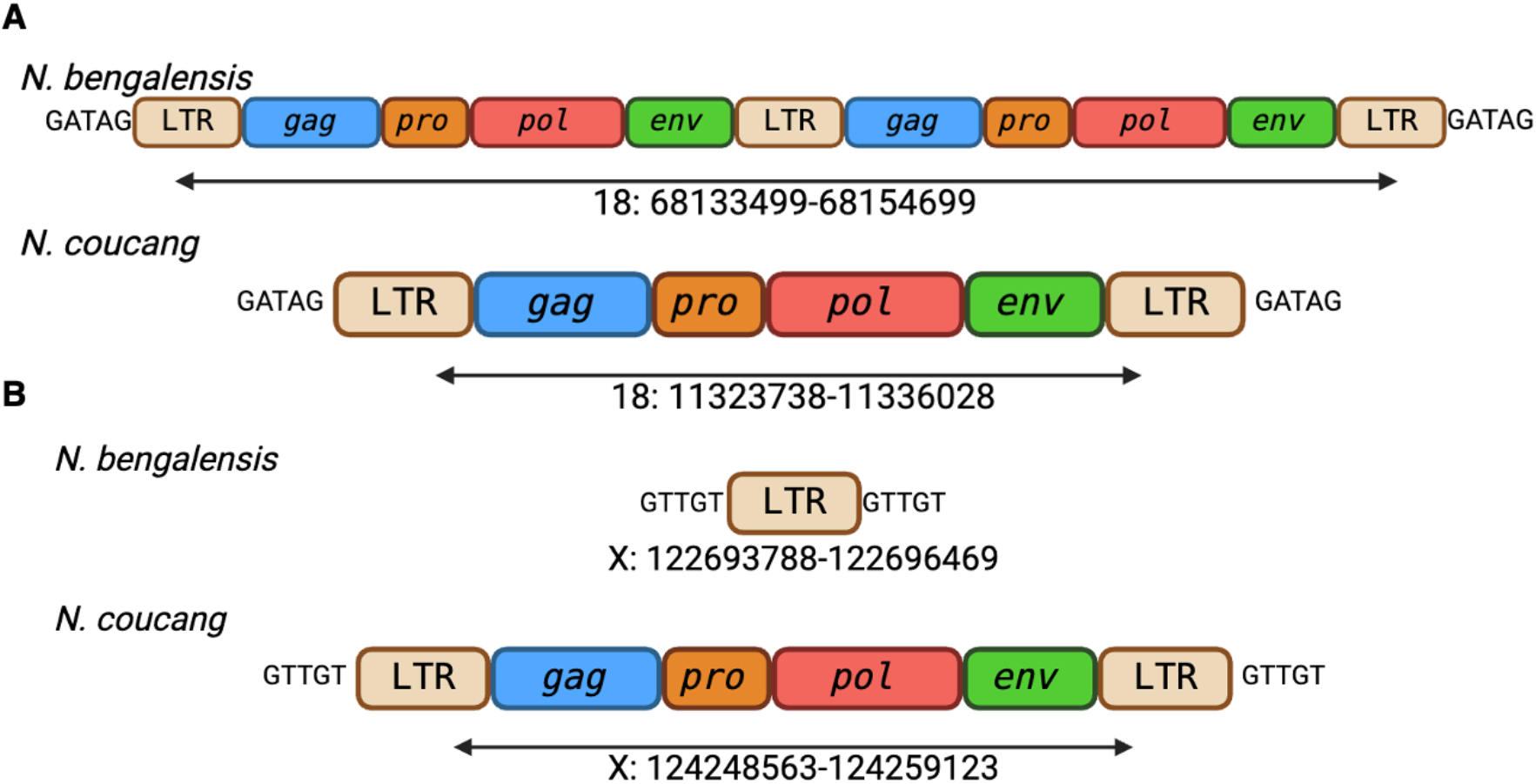
Graphical representation of examples of genomic events of LERV1 between *N. bengalensis and N. coucang*. (A) represents where there has been a segmental duplication creating a triplet LTR with two internal sequences. (B) represents an example of homologous recombination, where in *N. bengalensis* the ERV underwent homologous recombination but did not in *N. coucang*. Letters at start and end of LTR are the TSDs.

### 3.4 Dating of LERV1 insertions

As the LTRs are identical at the time of insertion, the divergence between the 5’ and 3’ LTR can be used to estimate the age of the insertion. Based on the divergence of the LTRs in complete full length insertions, the youngest LERV1 insertion is approximately 1.7 million years old and oldest may be ∼63 million years old (Table 1; Figure 5).

**Table 1:** LTR similarity of complete full-length insertions within *N. bengalensis* and *N. coucang* within a genome and between genomes in 5’ and 3’ LTR. Sorted here by estimated age based on *N. coucang*. Estimated age of insertion based on *N. coucang* except for ERV marked with * as they were solo LTRs in *N. coucang*. Multiple insertions could not be found in *N. bengalensis* and have been marked with ‘Region not found’ as searching for the flanking region of the ERV revealed no results.

| Full Length ERV in *N. coucang* | Full length ERV in *N. bengalensis* | 5' and 3' LTR similarity in *N. coucang* (%) | 5' and 3' LTR similarity in *N. bengalensis* (%) | Similarity of 5' LTR between *N. coucang* and *N. bengalensis* | Similarity of 3' LTR between *N. coucang* and *N. bengalensis* | Estimated age of insertion |
| --- | --- | --- | --- | --- | --- | --- |
| X: 124248563-124259123 | Solo | 99.7 |  |  |  | 1772546.8 |
| 2: 47031798-47039527 | 2: 137193135-137202130 | 99.3 | 95.8 | 96.7 | 99.6 | 4300261.5 |
| 20: 24900076-24909605 | 20: 41437738-41446756 | 99.3 | 97.9 | 99.3 | 98.2 | 4300261.5 |
| 14: 62267142-62277804 | 5' Truncated | 99.2 |  |  | 95.2 | 4431367.0 |
| 21: 33057945-33068610 | 21: 34962947-34972099 | 98.8 | 98.6 | 99.7 | 100 | 7068636.5 |
| X: 103734398-103744217 | Region not found | 98.8 |  |  |  | 7090187.2 |
| 5: 53187431-53194957 | Region not found | 98.5 |  |  |  | 8862734.0 |
| 20: 310325-320042 | Region not found | 98.4 |  |  |  | 9070130.3 |
| 20: 28209-38862 | Region not found | 98.4 |  |  |  | 9070130.3 |
| 8: 42690416-42705016 | 8: 43109200-43125182 | 98.1 | 94.0 | 95.5 | 100 | 11091803.2 |
| 19: 34353886-34364244 | 3' Truncated | 98.1 |  | 95.0 |  | 11180679.8 |
| 19: 37620478-37631360 | 19: 37288119-37297096 | 98.1 | 96 | 98.7 | 99.1 | 11197243.8 |
| 10: 28063825-28071582 | 10: 30226445-30235570 | 98.1 | 97.6 | 99.4 | 99.6 | 11331543.5 |
| 18: 11323738-11336028 | 18: 68133499-68154699 | 97.9 | 94.3 | 96.7 | 97.6 | 12040732.1 |
| 2: 144643661-144650652 | 2: 40398135-40406646 | 97.8 | 97.1 | 99.9 | 99.2 | 13028467.2 |
| 9: 127012398-127019684 | 9: 6944207-6952943 | 97.8 | 97.8 | 99.4 | 99.1 | 13055284.8 |
| 9: 53096813-53104969 | 9: 81375787-81384048 | 97.6 | 95.7 | 97.2 | 99.6 | 14039376.3 |
| 2: 138438690-138446251 | 2: 46370187-46379268 | 97.4 | 97.4 | 100 | 100 | 15363715.8 |
| Solo | 23: 8668601-8676540 |  | 97.3 |  |  | 15614093.1 |
| 4: 52113443-52128907 | 4: 90362357-90379212 | 97.3 | 97.2 | 99.9 | 100 | 15642924.2 |
| 6: 109789046-109804416 | 6: 97178488-97194222 | 97.2 | 96.6 | 99.8 | 99.6 | 16219183.6 |
| 22: 37965930-37977055 | Region not found | 97.2 |  |  |  | 16292790 |
| 22: 38113114-38122937 | Region not found | 97.1 |  |  |  | 17150305.3 |
| 5: 60467791-60474717 | Region not found | 96.9 |  |  |  | 18087855.3 |
| 5: 60066646-60074917 | Region not found | 96.9 |  |  |  | 18087855.3 |
| 20: 31905584-31915179 | 20: 36004074-36013094 | 96.8 | 96.7 | 100 | 99.2 | 18349687.1 |
| 15: 43375057-43383068 | 15: 53866152-53874258 | 96.6 | 96.8 | 100 | 99.9 | 19597596.0 |
| 1: 128253477-128261561 | 1: 128032638-128040828 | 96.6 | 96.3 | 99.8 | 99.1 | 19792182.1 |
| 8: 12617631-12626019 | 8: 12291873-12303285,<br>12636439-12644846 | 95.7 | 83.0 | 94.9 | 93.2 | 24224806.2 |
| 15: 49264183-49273173 | 15: 47917323-47926411 | 95.8 | 96.1 | 99.9 | 99.9 | 24428376 |
| 20: 31666264-31677164 | 20: 36230920-36241279 | 95.0 | 95.1 | 98.5 | 98.5 | 29155519 |
| 2: 106695258-106708893 | 2: 77805414-77819310 | 94.8 | 93.5 | 97.6 | 98.7 | 30191153.1 |
| X: 120489174-120506031 | X: 119049577-119065816 | 94.8 | 94.6 | 99.8 | 100 | 30496202.7 |
| 3: 63046647-63054274 | 3: 99111834-99120909 | 94.5 | 94.5 | 100 | 100 | 32076984.8 |
| X: 102141461-102151973 | X: 100642818-100652852 | 94.2 | 94.2 | 100 | 100 | 33479226.5 |
| 1: 180409850-180418816 | 1: 180687385-180696471 | 94.2 | 94.2 | 100 | 100 | 33752995.6 |
| 13: 38458101-38465647 | 13: 37318128-37327177 | 93.8 | 93.7 | 99.7 | 99.6 | 35928926.1 |
| 1: 34886678-34897666 | 3' Truncated | 93.8 |  | 98.0 |  | 36080952.5 |
| 10: 127504182-127513221 | 10: 122531696-122540844 | 93.7 | 93.8 | 100 | 99.9 | 36693956.1 |
| X: 94580487-94590236 | X: 93078643-93087724 | 93.1 | 93.9 | 100 | 99.9 | 40181060.8 |
| 20: 32544081-32554130 | 20: 35384555-35392095 | 92.9 | 88.9 | 94.7 | 99.7 | 40629957.82 |
| 20: 29859237-29877567 | 20: 37397166-37414564 | 93.0 | 93.0 | 100 | 100 | 40990463.4 |
| Solo | 5: 97346786-97356098 |  | 92.2 |  |  | 45069406.9 |
| X: 73374168-73385924 | Region not found | 91.9 |  |  |  | 47228193.6 |
| 7: 136140584-136148449 | 7: 1572227-1581451 | 91.8 | 91.8 | 100 | 100 | 47482378.8 |
| X: 65512009-65535211 | X: 64530215-64532926,<br>64552279-64554991 | 90.7 | 90.7 | 100 | 100 | 54197871.5 |
| X: 102642308-102661464 | X: 101153588-101170718 | 89.1 | 89.0 | 100 | 99.9 | 63230850.1 |

**Figure 5.**
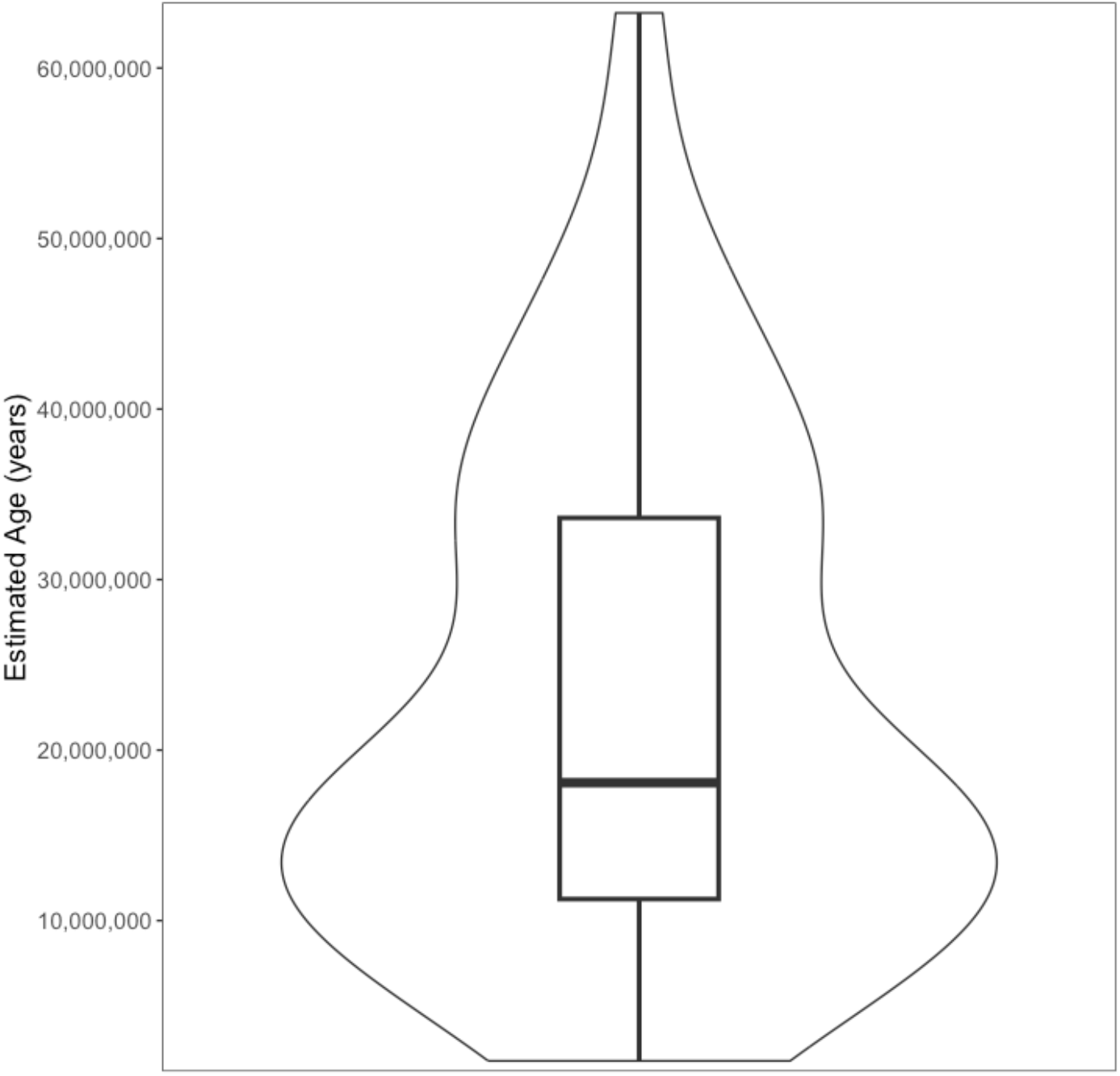
Range of estimated age of viral insertions of LERV1 based on 5’ and 3’ LTR divergences.

Insertions at the same locus in *N. bengalensis* and *N. coucang* would indicate the insertion occurring prior to the divergence of these two species. There was high sequence similarity between full-length insertions, and the truncated LTRs (Supplementary Table 3), with the majority of these shared insertions indicating >99% homology, and a few insertions were 100% identical in the two species. Given that these species are thought to have diverged approximately 5 MYA (Blair et al., 2023), this was surprising. For loci that inserted before the divergence of these two species we would expect the 5’ and 3’ LTRs of those full length elements to be more similar within a species than between. Focusing on the complete full length elements that were present in both species we found the differences between 5’ and 3’ LTRs between species was less than that within a species (Table 1).

Due to the surprisingly high similarity between insertions and LTRs between the *N. bengalensis* and *N. coucang* genomes, we compared mtDNA to confirm the classification of these species. A number of mtDNA genes are available on Genbank for these two species (Blair et al., 2023; Pozzi et al., 2015). Cytochrome B (cytb), NADH dehydrogenase subunit 4 (ND4) and cytochrome c oxidase subunit I (COX1) sequences of *N. bengalensis, N. coucang, and N. javanicus* (Supplementary Table 4) were downloaded from Genbank .We extracted the same genes from the reference genomes for *N. bengalensis* and *N. coucang* and constructed a Bayesian phylogeny to confirm the species identification. The mitochondrial sequences from both the reference genome and reference mitochondria (NC_002765) of *N. coucang* are placed within the *N. bengalensis* with strong support across all genes. (Figure 6).

**Figure 6.**
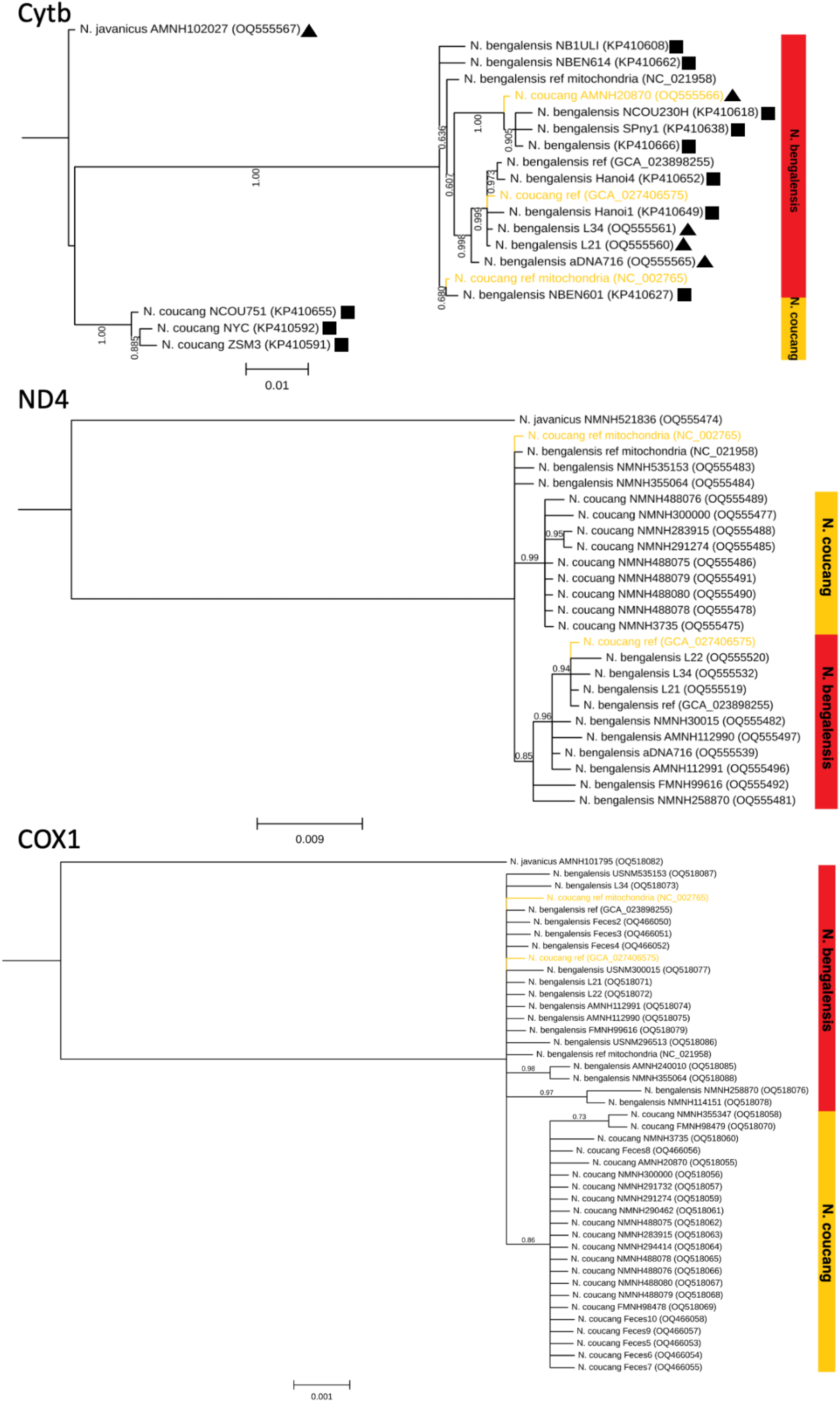
Phylogenetic tree of three mitochondrial genes Cytb, ND4 and COXI of slow lorises inferred from Bayesian (MrBayes) analysis with posterior probability shown. Scale bar represents substitutions per site. In Cytb triangle represents from Blair et al., (2023) and square represents from Pozzi et al., (2015). Tips in yellow represent sequences which come from individuals which have previously been identified as *N. coucang*.

### 3.5 Identification of novel full-length ERV in other primates

BLASTn searches of the consensus 3’ LTR and *env* identified the novel full-length ERV in three other species reference genomes; *X. pygmaeus, L. tardigradus* and *L. lydekkerianus*. No hits with substantial or any query coverage or percentage identity were found in *P. potto, O. garnetti, L. catta* or *H. sapiens*.

## 4 Discussion

A substantial number of ERVs have been identified in great apes and other Haplorhini, while comparatively fewer studies have focused on Strepsirrhini, particularly lorises. This study represents the first detailed examination of ERVs specifically in slow lorises within the genus *Nycticebus*, and the first to identify all major retroviral genes. Using the LTR of an incomplete (*gag, env* missing) “Unknown HERV” insertion reported in a previous study (Li et al., 2022), we identified 47 full length viral insertions (Table 1) which had remnants of all genes across two genomes of *Nycticebus*, however none of the genes were intact, suggesting that this is not a recent genome invasion of this family of retrovirus Over 100 insertions were incomplete copies (missing internal genes, 5’ or 3’ truncated, see Supplementary Table 3). The BLAST results indicate numerous solo LTRs (∼1400) present in the *Nycticebus* genome, which is similar to observations of other primate ERVs, where the solo LTRs are ten times more numerous than the full length insertions (Belshaw et al., 2007).

Phylogenetic analysis reveals this virus is a betaretrovirus, most closely related and most similar to HERV-K across all genes (Figure 2). Despite being most similar to HERV-K and the LTR being previously labelled as being from an “Unknown HERV” this virus does not appear to be a HERV and is unique to Asian lorises. This relationship may warrant further investigation as throughout the phylogenetic tree retroviruses appear to approximately cluster with retroviruses from similar or closely related species, but LERV1 is closest to HERV-K. The novel retrovirus identified here is distinct from the RV Slow Loris (Gifford et al., 2005) previously identified within the *Nycticebus* genus.

The HERV-K betaretrovirus is one of the most well studied family of retroviruses as it is the most recently integrated endogenous retrovirus in the human genome, with some loci being insertionally polymorphic (Hohn et al., 2013). HERV-K has had a significant impact in the human genome, with certain loci implicated in addiction (Karamitros et al., 2018), in addition to the numerous links between HERV-K and diseases such as various cancers (Dervan et al., 2021; Rivas et al., 2022), and neurodegenerative disorders (Adler et al., 2024). With comparable numbers of full length proviruses and solo LTRs, it is likely that LERV1 has had a similar impact in the genome evolution of the subfamily Lorisinae. Estimates of the age of the HERV-K family are also similar to our age estimates of LERV1 calculated here, ranging from 1.7 MYA to 63 MYA (Table 1), providing further support for the idea that LERV1 has had a significant impact on genome evolution in these species. However, the genomes of the smaller primates are not as well annotated as the larger primate genomes to investigate this currently. The virus could be identified exclusively in the subfamily Lorisinae suggesting that the oldest insertion should have occurred approximately 30Mya in the common ancestor of the subfamily before the divergence into the genera *Loris, Nycticebus* and *Xanthonycticebus* (Pozzi et al., 2015); our age estimates for some loci exceed this date, which could indicate that gene conversion has occurred (Jedlicka et al., 2020), in which case these estimates are unlikely to be accurate. The subfamily Lorisinae is found exclusively in Asia whilst its sister subfamily Perodicticinae is in Africa, indicating that the initial LERV1 integration likely happened somewhere within Asia (specifically India) after the invasion of the common ancestor of Lorisinae colonized Asia (Ali & Aitchison, 2008).

We observed several instances of insertional polymorphism of LERV1 loci in the *N. bengalensis* and *N. coucang* genome (Table 1, Supplementary Table 3). Given the divergence times of these two species are estimated to be between 4 and 8 MYA (Blair et al., 2023; Pozzi et al., 2015), this would suggest that these particular loci inserted in the common ancestor at around this time.

However, our calculations of insertion times for three loci which exhibit insertional polymorphism through homologous recombination events based on the LTR divergence of the full length provirus, range from 1.8 MYA to 45 MYA. For the youngest of these (X:124248563-124259123 in *N. coucang*), the full-length insertion in *N. coucang* had LTRs that are 99.7% similar, and the solo LTR within *N. bengalensis* is 99.24% to 99.54% similar to the 5’ and 3’ LTRs in *N. coucang* respectively; looking at the remaining proviruses, in the large majority of loci, comparison of the LTRs of each locus indicated a higher degree of similarity between the 5’ and 3’ LTRs of the two species, as opposed to within the species (Table 1). This was unexpected given the divergence time of *N. bengalensis* and *N. coucang*, and the range of proviral integration times. Due to this very high level of similarity a brief examination of the mitochondrial genes from the *N. bengalensis* and *N. coucang* reference genomes was undertaken to determine if they are different species. Our results show that the current reference genome of *N. coucang* is not a pure *N. coucang*, and is either a hybrid whose mother was a *N. bengalensis*, or a *N. bengalensis* from a different population than the reference genome of *N. bengalensis* (Figure 6). We also show that the reference mitochondrial *N. coucang* (NC_002765) is also unlikely to have come from *N. coucang*. Examination of nuclear DNA will be required to establish the true nature of these genomes and whether they are hybrids. As the *N. coucang* reference genome comes from an individual born in San Diego Zoo in 1989 (https://www.ncbi.nlm.nih.gov/biosample/SAMN28408555/) it is possible that the parents may have been misidentified (where one or both were *N. bengalensis*), as in the past it has been challenging to differentiate between the two species, and there have been species reclassifications (Nekaris, 2014; Nekaris & Starr, 2015; Pozzi et al., 2015). Regardless, the similarity of viral elements between species indicates a more recent common ancestor of the *N. bengalensis* and *N. coucang* reference genomes compared to the expected ∼4 to 8 MYA species divergence.

The discovery of this retrovirus within the slow lorises provides insights into both the genome evolution of these species and viruses themselves. Through the examination of this virus, we have found multiple genomic events including homologous recombination, duplication and inversions. Further examination could be carried out to discover if any solo LTRs are close to genes and therefore may be acting as a promoter or enhancer (e.g. Pi et al., 2004). Examination of this virus allowed us to determine that it is likely that the currently published reference genome of *N. coucang* is incorrect and may be a hybrid with *N. bengalensis*, or a *N. bengalensis* from a different population. Both *N. bengalensis* and *N. coucang* are endangered (Nekaris, Al-Razi, et al., 2020; Nekaris, Poindexter, et al., 2020), and therefore it is crucial to correctly determine the classification of these species and genomes.

## Notes

### Competing Interest Statement

The authors have declared no competing interest.

