## Supplementary Information for "Novel Endogenous Retrovirus in the Slow Loris"

TGTAGGGTAATATTTAGAAATAAACATATCCTACATCAAGCACACAAAGTGAAC
TCAGTGTGAGTTTAAAATGACTTGCTTTTCTTTCAGAGGTTTAAGGGACATGGGC
GCCCCTGGGATACACACAGAATTGGTTTACCTGGAGTTATAAATTCCATAAGCCA
TTTCCCTGAAGCTGTACACCCCATGAAGCCTGAGATCAAAGGGCATGCTGCCCTT
CCTCAGAGATAGGGGCCAGCGGGTTGATGTATGTTAACAACATATTGTCAGAGA
TCTATATGTATTCTGCCCTGAGGGAAAGACACAGTCATTACTTCATACTGAGTAA
ACTGAGTGAACGCTTGAAGATATTCTTCAATTAACTCTGTTCTCCTATGGCTACTT
TCTTCTTTGTGATCTTTATTTAGATACCTGCTCAGAAATTAGTCAGTGTTTAGAAG
AAGAGGTTTACTACAAAAGTGATTGTATTGATATGATAACAAAATATTTCCTTTC
AAGTTAATTTCAGATAGGTAAAATAAGATAATGATGGAACTAACAGATGATAGA
TCAAGTGTGTACATAACTAAAAAAGTATAAAATCAAAACCCTGAATAAAGCCTT
TCTCTACACACCATCCCATCTCGTGTAGTCTGTTTCTTGACTTATTAATTACAGGA
GTGAGTTTACACCCACGGCAAAGCTGCTGTTTGTTTGTGTCTCCTGTCACGCAGC
A

**Supplementary Figure 1: LTR of an “Unknown HERV” from Li et al., (2022) used in BLASTn screening of the loris genomes.**

**Supplementary Table 1: Sequences used in phylogenetic analysis of SLERV-B-1. N/A indicates not included in analysis for the gene.**

| Name | Accession number or reference | Gag genera | Pro genera | Pol genera | Env genera |
|---|---|---|---|---|---|
| Avian leukaemia virus | NC_015116 | Alpha | Alpha | Alpha | Alpha |
| Avian leukosis virus | KU375453 | Alpha | Alpha | Alpha | Alpha |
| Black Syrian hamster retrovirus | MK304634 | Beta | Beta | Beta | Beta |
| Bovine retrovirus CH15 | KU720628 | Beta | Beta | Beta | Beta |
| Enzootic nasal tumour virus 2 | MK164396 | Beta | Beta | Beta | Beta |
| HERV-K113 | NC_022518 | Beta | Beta | Beta | Beta |
| HERV-K115 | AY037929 | Beta | Beta | Beta | Beta |
| Jaagsiekte sheep retrovirus | NC_001494 | Beta | Beta | Beta | Beta |
| Mouse mammary tumor virus | NC_001503 | Beta | Beta | Beta | Beta |
| Ovine enzootic nasal tumour virus | NC_007015 | Beta | Beta | Beta | Beta |
| Desmodus rotundus ERV | NC_027117 | Beta | Beta | Beta | Type-D Beta |
| Mason-Pfizer monkey virus | NC_001550 | Beta | Beta | Beta | Type-D Beta |
| Prosimian retrovirus 1 | MT787217 | Beta | Beta | Beta | Type-D Beta |
| Python molurus endogenous retrovirus | AF500296 | Beta | Beta | Beta | Type-D Beta |
| Simian endogenous retrovirus | U85505 | Beta | Beta | Beta | Type-D Beta |
| Simian retrovirus 1 | M11841 | Beta | Beta | Beta | Type-D Beta |
| Simian retrovirus 2 | MF805808 | Beta | Beta | Beta | Type-D Beta |
| Simian retrovirus 4 | NC_014474 | Beta | Beta | Beta | Type-D Beta |
| Simian retrovirus 8 | NC_031326 | Beta | Beta | Beta | Type-D Beta |
| Squirrel monkey retrovirus | NC_001514 | Beta | Beta | Beta | Type-D Beta |

| | | | | | |
|---|---|---|---|---|---|
| Cervid endogenous betaretrovirus 1 | OL547611.1 | Beta | Beta | Beta | N/A |
| RV_Slow_Loris | Gifford et al. (2005) | N/A | N/A | Beta | N/A |
| Human T-lymphotropic virus 2 | NC_001488 | Delta | Delta | Delta | Delta |
| Simian T-cell lymphotropic | NC_011546 | Delta | Delta | Delta | Delta |
| Baboon endogenous virus | NC_022517 | N/A | Gamma | Gamma | Gamma |
| Gibbon ape leukaemia virus | NC_001885 | N/A | Gamma | Gamma | Gamma |
| Grey mouse lemur ERV-Fc | Diehl et al. (2016) | N/A | N/A | Gamma | Gamma |
| Koala retrovirus | NC_039228 | N/A | Gamma | Gamma | Gamma |
| Woolly monkey sarcoma virus | KT724051 | N/A | Gamma | Gamma | Gamma |
| Brown greater galago prosimian foamy virus | NC_039023 | N/A | Spuma | Spuma | N/A |
| Rhesus macaque simian foamy virus | NC_039238 | N/A | Spuma | Spuma | N/A |

**Supplementary Table 2: Chromosomes of *N. coucang* and inferred chromosomes of *N. bengalensis* based on BLASTn and viral insertions.**

| ***N. coucang* chromosome** | **Genbank ID** | **Size (bp)** | ***N. bengalensis* chromosome** | **Genbank ID** | **Size (bp)** | **Inferred Chromosome** |
|---|---|---|---|---|---|---|
| 1 | CM050137.1 | 196,249,132 | LG01 | CM043610.1 | 196,305,487 | 1 |
| 2 | CM050138.1 | 184,820,795 | LG02 | CM043611.1 | 185,133,867 | X |
| 3 | CM050139.1 | 161,203,717 | LG03 | CM043612.1 | 184,494,762 | 2 |
| 4 | CM050140.1 | 148,538,759 | LG04 | CM043613.1 | 160,225,771 | 3 |
| 5 | CM050141.1 | 142,425,405 | LG05 | CM043614.1 | 142,930,404 | 4 |
| 6 | CM050142.1 | 138,330,575 | LG06 | CM043615.1 | 140,782,244 | 5 |
| 7 | CM050143.1 | 137,715,312 | LG07 | CM043616.1 | 132,450,243 | 7 |
| 8 | CM050144.1 | 136,588,019 | LG08 | CM043617.1 | 130,794,859 | 9 |
| 9 | CM050145.1 | 135,917,784 | LG09 | CM043618.1 | 128,886,875 | 11 |
| 10 | CM050146.1 | 133,885,751 | LG10 | CM043619.1 | 128,767,214 | 8 |

| | | | | | | |
|---|---|---|---|---|---|---|
| 11 | CM050147.1 | 128,238,841 | LG11 | CM043620.1 | 128,728,782 | 10 |
| 12 | CM050148.1 | 112,708,720 | LG12 | CM043621.1 | 126,158,910 | 6 |
| 13 | CM050149.1 | 103,192,619 | LG13 | CM043622.1 | 108,846,978 | 12 |
| 14 | CM050150.1 | 99,988,313 | LG14 | CM043623.1 | 98,296,284 | 14 |
| 15 | CM050151.1 | 97,885,895 | LG15 | CM043624.1 | 96,198,676 | 13 |
| 16 | CM050152.1 | 96,499,272 | LG16 | CM043625.1 | 95,554,990 | 16 |
| 17 | CM050153.1 | 91,048,355 | LG17 | CM043626.1 | 93,099,192 | 15 |
| 18 | CM050154.1 | 80,579,731 | LG18 | CM043627.1 | 90,664,049 | 17 |
| 19 | CM050155.1 | 72,490,086 | LG19 | CM043628.1 | 72,079,205 | 19 |
| 20 | CM050156.1 | 69,287,667 | LG20 | CM043629.1 | 70,248,661 | 18 |
| 21 | CM050157.1 | 67,625,389 | LG21 | CM043630.1 | 62,093,175 | 20 |
| 22 | CM050158.1 | 61,455,708 | LG22 | CM043631.1 | 61,817,876 | 22 |
| 23 | CM050159.1 | 38,949,463 | LG23 | CM043632.1 | 56,716,581 | 21 |
| 24 | CM050160.1 | 33,060,932 | LG24 | CM043633.1 | 31,240,522 | 24 |
| X | CM050161.1 | 187,328,519 | LG25 | CM043634.1 | 16,045,275 | 23 |
| Y | CM050162.1 | 35,646,755 | | | | |

**Supplementary Table 3: Full length elements and truncated SLERV-B-1 elements in *N. coucang* and their equivalent in *N. bengalensis* based on matching TSDs and flanking regions. Similarity between species is calculated as pairwise identity across the whole insertion. CFL = complete full length. Sorted here by chromosome. Orientation is only listed for *N. coucang* as it is believed to be the higher quality genome. * indicates elements with large indels removed**

| **Chromosome in *N. coucang*** | **Co-ordinates in *N. coucang*** | **Orientation in *N. coucang*** | **State in *N. coucang*** | **Chromosome in *N. bengalensis*** | **Co-ordinates in *N. bengalensis*** | **State in *N. bengalensis*** | **Predicted genomic event within *N. bengalensis*** | **Predicted genomic event within *N. coucang*** | **Similarity between species (%)** |
|---|---|---|---|---|---|---|---|---|---|
| 1 | 46507044-46510269 | - | gag, env | 1 | 46232953-46235659 | 3’ Truncated | Truncation | | 100 |
| 1 | 34887678-34896666 | + | CFL | 1 | 34474841-34477547 | 3’ Truncated | Truncation | | 88.738 |
| 1 | 57200495-57206971 | + | gag, pro, env | 1 | 56990722-56997198 | gag, pro, env | | | 99.383 |
| 1 | 128253477-128261561 | + | CFL | 1 | 128032638-128040828 | CFL | | | 99.245 |
| 1 | 180409850-180418816 | + | CFL | 1 | 180687385-180696471 | CFL | | | 99.605 |
| 1 | 62712062-62714764 | - | 5' Truncated | 1 | 62558245-62560947 | 5' Truncated | | | 99.926 |
| 1 | 175390395-175393108 | - | 3' Truncated | 1 | 175635952-175638665 | 3' Truncated | | | 100 |
| 1 | 86804063-86806781 | + | 3' Truncated | 1 | 86718281-86720999 | 3' Truncated | | | 98.566 |
| 2 | 47031798-47039527 | - | CFL | 2 | 137193135-137202130 | CFL | | | 98.761 |
| 2 | 138438690-138446251 | - | CFL | 2 | 46370187-46379268 | CFL | | | 99.835 |
| 2 | 144643661-144650652 | - | CFL | 2 | 40398135-40406646 | CFL | | | 99.847 |

| | | | | | | | | | |
|---|---|---|---|---|---|---|---|---|---|
| 2 | 2441337-2448751 | + | gag, pro, env | 2 | 182162388-182171857 | Triplet (LTR-LTR-gag, pro, env – LTR) | Duplication of LTR | | 89.171 |
| 2 | 34655973-34662570 | + | pol, env | 2 | 149679446-149686170 | pol, env | | | 99.941 |
| 2 | 106695258-106708893 | + | CFL | 2 | 77805414-77819310 | CFL | | | 98.73 |
| 2 | 1504475-1507184 | - | 5' Truncated | 2 | 183095608-183098317 | 5' Truncated | | | 99.594 |
| 2 | 95974860-95977576 | + | 5' Truncated | 2 | 87970550-87973265 | 5' Truncated | | | 98.971 |
| 2 | 143444189-143446909 | + | 3' Truncated | 2 | 41536054-41538774 | 3' Truncated | | | 99.963 |
| 3 | 63046647-63054274 | - | CFL | 3 | 99111834-99120909 | CFL | | | 99.945 |
| 3 | 26505737-26508436 | - | 5' Truncated | 3 | 133621016-133623715 | 5' Truncated | | | 98.626 |
| 3 | 63654279-63656974 | + | 5' Truncated | 3 | 98500148-98502843 | 5' Truncated | | | 100 |
| 4 | 135430104-135433433 | - | gag, env | 4 | 6042451-6045176, 6091360-6094080 | 3’ Truncated and 5’ Truncated | Truncation and Inversion of an LTR | Duplication and inversion (4: 135430104-135433433, 4: 136857158-136861849, 4: 137061734-137066425) | 99.451 & 88.611 |
| 4 | 52113443-52128907 | + | CFL | 4 | 90362357-90379212 | CFL | | | 84.362 |

| | | | | | | | | | |
|---|---|---|---|---|---|---|---|---|---|
| 4 | 136857158-136861849 | + | gag, env | 4 | 6042451-6045176, 6091360-6094080 | 3' Truncated and 5' Truncated | Truncation and Inversion of an LTR | Duplication and inversion (4: 135430104-135433433, 4: 136857158-136861849, 4: 137061734-137066425) | 99.451 & 88.611 |
| 4 | 137061734-137066425 | + | gag, env | 4 | 6042451-6045176, 6091360-6094080 | 3' Truncated and 5' Truncated | Truncation and Inversion of an LTR | Duplication and inversion (4: 135430104-135433433, 4: 136857158-136861849, 4: 137061734-137066425) | 99.451 & 88.611 |
| 4 | 87656594-87659290 | - | 3' Truncated | 4 | 54398993-54401701 | 3' Truncated | | | 95.622 |
| 4 | 145404499-145407208 | - | 5' Truncated | 4 | 3716613-3719321 | 5' Truncated | | | 99.889 |
| 5 | 53187431-53194957 | - | CFL | 5 | | Region not found | | | |
| 5 | 60467791-60474717 | - | CFL | 5 | | Region not found | | Duplication and inversion (5: 60467791-60474717, 5: 60066646-60074917) | |

| | | | | | | | | | |
|---|---|---|---|---|---|---|---|---|---|
| 5 | 93701941-93704494 | - | gag, env | 5 | 48765554-48769633 | gag, env | | | 99.559 |
| 5 | 93815050-93827133 | - | Two Solo | 5 | 48645172-48658752 | Two Solo | | | 99.941 |
| 5 | 125614825-125616664 | - | Two Solo | 5 | 16588467-16591587 | Two Solo | | | 98.341 |
| 5 | 38309066-38322487 | + | Two Solo | 5 | 102289321-102302839 | Two Solo | | | 99.741 |
| 5 | 60066646-60074917 | + | CFL | 5 | | Region not found | | Duplication and inversion (5: 60467791-60474717, 5: 60066646-60074917) | |
| 5 | 43189280-43191935 | + | Solo | 5 | 97346786-97356098 | CFL | | Homologous recombinatio n | 90.137 |
| 5 | 35207252-35209976 | + | 5' Truncated | 5 | 105397218-105399942 | 5' Truncated | | | 99.509 |
| 6 | 109789046-109804416 | + | CFL | 6 | 97178488-97194222 | CFL | | | 99.043 |

| | | | | | | | | | |
|---|---|---|---|---|---|---|---|---|---|
| 6 | 89516673-89519393, 91064763-91067483, 96689056-96691776, 97905997-97908717, 82291713-82294433, 82852106-82854824, 88944676-88947396 | -, -, -, -, +, +, + | 5' Truncated | 6 | 85523445-85526165 | 5’ Truncated | | Duplications and inversion | 91.102 - 95.775 |
| 7 | 136140584-136148449 | - | CFL | 7 | 1572227-1581451 | CFL | | | 99.566 |
| 7 | 31356172-31363182 | + | gag, env | 7 | 101016669-101023760 | gag, env | | | 99.087 |
| 7 | 32466046-32472425 | + | gag, pro, env | 7 | 99904465-99911169 | gag, pro, env | | | 99.33 |
| 7 | 50776817-50786935 | + | Two Solo | 7 | 87489776-87499991 | Two Solo | | | 99.394 |
| 7 | 64498461-64517619 | + | Two Solo | 7 | 73742341-73761635 | Two Solo | | | 99.679 |
| 8 | 42690416-42705016 | - | CFL | 8 | 43109200-43125182 | CFL | Inversion (in either) | Inversion (in either) | 97.03 |
| 8 | 12617631-12626019 | + | CFL | 8 | 12291873-12303285, 12636439-12644846 | CFL | Duplication with insertion (12291873-12303285) | | 89.415 & 92.26 |
| 9 | 51512182-51519035 | - | Two Solo | 9 | 82887572-82895915 | Two Solo | | | 99.868 |

| | | | | | | | | | |
|---|---|---|---|---|---|---|---|---|---|
| 9 | 52014261-52018097 | - | gag, env | 9 | 82397565-82407452 | Triplet (LTR – gag, env – LTR – gag, env – LTR) | Duplication | | 98.397 & 97.912 |
| 9 | 52018791-52022212 | - | Two Solo | 9 | 82393433-82398383 | Two Solo | | | 99.867 |
| 9 | 127012398-127019684 | - | CFL | 9 | 6944207-6952943 | CFL | | | 98.17 |
| 9 | 53096813-53104969 | + | CFL | 9 | 81375787-81384048 | CFL | | | 99.129 |
| 9 | 51189498-51192208 | - | 5' Truncated | 9 | 83219367-83222078 | 5' Truncated | | | 99.189 |
| 10 | 28049430-28063338 | - | Two Solo | 10 | 30234780-30249973 | Two Solo | | | 99.724 |
| 10 | 28063825-28071582 | - | CFL | 10 | 30226445-30235570 | CFL | | | 94.636 |
| 10 | 29284048-29292578 | - | gag, pro, env | 10 | 28986606-28989289 | 3’ Truncated | Truncation | | 99.052 |
| 10 | 80075455-80078436 | - | Two Solo | 10 | | Region not found | | | |
| 10 | 88899555-88905446 | + | gag, pro, env | 10 | 84065323-84071121 | gag, env | Deletion of pro | | 90.235 |
| 10 | 127504182-127513221 | + | CFL | 10 | 122531696-122540844 | CFL | | | 99.956 |
| 10 | 51436069-51438776 | - | 5' Truncated | 10 | 48640044-48642751 | 5' Truncated | | | 99.647 |
| 10 | 34407121-34409837 | + | 5' Truncated | 10 | 31039813-31042529 | 5' Truncated | Inversion (in either) | Inversion (in either) | 99.742 |
| 11 | 33572164-33589266 | - | Two Solo | 11 | 95096887-95115489 | Two Solo | | | 99.93 |
| 11 | 50979880-50987230 | - | Two Solo | 11 | 77460446-77469355 | Two Solo | | | 99.102 |

| | | | | | | | | | |
|---|---|---|---|---|---|---|---|---|---|
| 11 | 95276552-95282006 | - | Two Solo | 11 | 33132878-33139691 | Two Solo | | | 97.965 |
| 11 | 105810380-105816635 | + | pro, pol, env | 11 | 22521325-22527916 | pro, pol, env | | | 93.843 |
| 11 | 34558888-34561579 | + | 5' Truncated | 11 | 94108556-94111247 | 5' Truncated | | | 99.963 |
| 11 | 41944382-41947101 | + | 5' Truncated | 11 | 86661956-86664675 | 5' Truncated | | | 99.779 |
| 11 | 83043956-83046667 | + | 5' Truncated | 11 | 45388131-45390842 | 5' Truncated | | | 99.41 |
| 12 | 75640065-75644175 | - | gag, pro, env | 12 | 37105592-37111215 | gag, pro, env | | | 98.846 |
| 12 | 17602846-17605559 | - | Solo | 12 | 91086085-91089009 | Two Solo | Duplication of LTR | | 100 |
| 12 | 96214909-96217571 | - | 5' Truncated | 12 | 16192225-16194913 | 5' Truncated | | | 99.812 |
| 12 | 10356712-10359426 | + | 5' Truncated | 12 | 98579715-98582429 | 5' Truncated | | | 98.566 |
| 12 | 69637507-69640217 | + | Solo | 12 | 43186666-43188212 | Two Solo | Duplication of LTR | | 97.977 |
| 13 | 1901377-1909806 | + | pro, pol, env | 13 | 1898260-1907130 | pro, pol, env | | | 96.879 |
| 13 | 43696114-43700735 | + | gag, env | 13 | 40414329-40419074 | gag, env | | | 99.621 |
| 13 | 19558224-19561418 | - | gag, env | 13 | 19586819-19591534 | gag, env | | | 99.915 |
| 13 | 38458101-38465647 | - | CFL | 13 | 37318128-37327177 | CFL | | | 99.757 |
| 13 | 43788005-43794599 | - | Two Solo | 13 | 40505400-40513538 | Two Solo | | | 99.779 |
| 13 | 2178156-2180864 | - | 5' Truncated | 13 | 2178784-2181492 | 5' Truncated | | | 100 |

| | | | | | | | | | |
|---|---|---|---|---|---|---|---|---|---|
| 13 | 43204429-43207147 | - | 3' Truncated | 13 | 39906419-39909137 | 3' Truncated | | | 99.963 |
| 13 | 1696266-1698976 | + | 5' Truncated | 13 | 1694166-1699235 | env | Duplication of LTR | | 99.416 |
| 13 | 45598917-45601637 | + | 3' Truncated | 13 | 42308800-42311520 | 3' Truncated | | | 99.816 |
| 13 | 74934052-74936766 | + | 5' Truncated | 13 | 67805905-67808619 | 5' Truncated | | | 99.963 |
| 14 | 62268142-62276804 | + | CFL | 14 | 60603920-60606604 | 5’ Truncated | Truncation | | 83.296 |
| 14 | 87298935-87304315 | + | gag, env | 14 | 85556873-85562401 | gag, env | | | 99.602 |
| 15 | 28623510-28630086 | + | Two Solo | 15 | 66882663-66889361 | Two Solo | | | 99.955 |
| 15 | 43375057-43383068 | + | CFL | 15 | 53866152-53874258 | CFL | | | 98.265 |
| 15 | 49264183-49273173 | + | CFL | 15 | 47917323-47926411 | CFL | | | 99.45 |
| 16 | 25041882-25055034 | - | Triplet (LTR - gag, pro, env – LTR – LTR) | 16 | 25215593-25228846 | Triplet (LTR - gag, pro, env – LTR – LTR) | Inversion (in either) | Inversion (in either) | 92.725 & 96.491 |
| 16 | 44508817-44511283 | - | Two Solo | 16 | 44088890-44091984 | Triplet (LTR - LTR - LTR) | Duplication of LTR (Middle LTR) | | 100 & 100 |
| 16 | 25400445-25419156 | - | pro, pol, env | 16 | 25571898-25579248 | pro, pol, env | | | 98.709 |
| 16 | 14570367-14573082 | - | Solo | 16 | 14663717-14665263 | Two Solo | Duplication of LTR | | 99.876 |

| | | | | | | | | | |
|---|---|---|---|---|---|---|---|---|---|
| 17 | 37801966-37804676 | - | Solo | 17 | 38157075-38159726 | Two Solo | Duplication of LTR | | 100 |
| 18 | 11323738-11336028 | + | CFL | 18 | 68133499-68154699 | Triplet (LTR – gag, pro, pol, env – LTR – gag, pro, pol, env – LTR) | Duplication | | 80.04 & 83.056 |
| 18 | 36712576-36719635 | + | gag, env | 18 | 44182215-44187422 | gag, env | | | 100 |
| 18 | 26848966-26851639 | - | 5' Truncated | 18 | 54047950-54050628 | 5' Truncated | | | 92.369 |
| 19 | 34353886-34364244 | + | CFL | 19 | 34764791-34767481 | 3’ Truncated | Truncation | | 94.299 |
| 19 | 37620478-37631360 | + | CFL | 19 | 37288119-37297096 | CFL | | | 99.566 |
| 19 | 38807390-38820765 | + | Two Solo | 19 | 38478797-38490353 | Two Solo | | | 99.836 |
| 19 | 56154383-56160494 | + | Two Solo | 19 | 55783005-55787295 | Two Solo | | | 98.441 |
| 19 | 8356972-8359685 | - | 5' Truncated | 19 | 8484857-8487570 | 5' Truncated | | | 100 |
| 19 | 35126697-35129373 | - | 3' Truncated | 19 | 35542350-35545026 | 3' Truncated | | | 98.73 |
| 19 | 35445791-35448497 | + | Solo | 19 | 35990950-35993435 | Two Solo | Duplication of LTR | | 96.605 |

| | | | | | | | | | |
|---|---|---|---|---|---|---|---|---|---|
| 20 | 310325-320042 | - | CFL | 20 | | Region not found | | Duplication and inversion (20: 310325-320042 & 20: 28209-38862) | |
| 20 | 24900076-24909605 | - | CFL | 20 | 41437738-41446756 | CFL | | | 98.649 |
| 20 | 29859237-29877567 | - | CFL | 20 | 37397166-37414564 | CFL | Deletion of zinc finger | | 97.556 |
| 20 | 31666264-31677164 | - | CFL | 20 | 36230920-36241279 | CFL | | | 98.114 |
| 20 | 31905584-31915179 | - | CFL | 20 | 36004074-36013094 | CFL | | | 99.126 |
| 20 | 32544081-32554130 | - | CFL | 20 | 35384555-35392095 | CFL | Large deletion in pol | | 76.149 |
| 20 | 28209-38862 | + | CFL | 20 | | Region not found | | Duplication and inversion (20: 310325-320042 & 20: 28209-38862) | |
| 20 | 34763018-34773346 | + | Two Solo | 20 | 34156644-34165109 | Two Solo | | | 98.106 |
| 20 | 33798621-33801334 | - | 5' Truncated | 20 | 34621291-34624004 | 5' Truncated | | | 99.926 |
| 20 | 61650274-61652987 | - | 5' Truncated | 20 | 7486165-7488882 | 5' Truncated | | | 99.669 |
| 20 | 13962777-13965487 | + | 5' Truncated | 20 | 49408867-49411576 | 5' Truncated | | | 98.414 |

| | | | | | | | | | |
|---|---|---|---|---|---|---|---|---|---|
| 20 | 33559927-33562640 | + | 3' Truncated | 20 | 34377487-34380200 | 3' Truncated | | | 99.889 |
| 21 | 11414152-11419347 | - | gag, env | 21 | 55689261-55693951 | gag, env | | | 98.367 |
| 21 | 16550188-16555925 | + | gag, pro | 21 | 50612540-50615212 | Solo | Homologous recombination | | 98.964 |
| 21 | 33057945-33068610 | + | CFL | 21 | 34962947-34972099 | CFL | | | 95.713 |
| 22 | 38113114-38122937 | - | CFL | 22 | | Region not found | | Duplication and inversion (22: 38113114-38122937, 22: 37965930-37977055) | |
| 22 | 37965930-37977055 | + | CFL | 22 | | Region not found | | Duplication and inversion (22: 38113114-38122937, 22: 37965930-37977055) | |
| 22 | 39586651-39589351 | + | 5' Truncated | 22 | 22180089-22182790 | 5' Truncated | | | 98.89 |

| | | | | | | | | | |
|---|---|---|---|---|---|---|---|---|---|
| Y | 11960026-11962727 | - | Solo | 23 | 136947-140987 | Two Solo | Probable misassembly and Duplication of LTR | | 89.347 |
| 23 | 9425876-9432504 | + | Solo | 23 | 7509697-7514518 | CFL | | Homologous recombination | 95.434 |
| 24 | 8653157-8655811 | + | gag, env | 24 | 8668601-8676540 | gag, env | | | 99.813 |
| X | 66267082-66274143 | - | gag, pro, env | X | 65289080-65295657 | gag, pro, env | | | 99.651 |
| X | 72614389-72620728 | - | gag, env | X | 71335356-71341177 | gag, env | | | 100 |
| X | 89812308-89819241 | - | gag, pro, env | X | 88355251-88367316 | Triplet (LTR - gag, pro, env – LTR – gag, pro, env) | Duplication | | 99.211 |
| X | 94580487-94590236 | - | CFL | X | 93078643-93087724 | CFL | | | 99.868 |
| X | 102141461-102151973 | - | CFL | X | 100642818-100652852 | CFL | | | 99.791 |
| X | 120489174-120506031 | - | CFL | X | 119049577-119065816 | CFL | | | 99.698 |
| X | 124159050-124164274 | - | env | X | 122632331-122636999 | env | | | 99.957 |

| | | | | | | | | | |
|---|---|---|---|---|---|---|---|---|---|
| X | 139533864-139545838 | - | gag, pro, env | X | 137497621-137509492 | gag, pro, env | | Duplication and inversion (X: 137497621-137509492 , X: 139222079-139235419) | 93.599 |
| X | 65512009-65535211 | + | CFL | X | 64530215-64532926, 64552279-64554991 | 3' Truncated and 5' Truncated | Large insertion in pol | | 92.308 |
| X | 73374168-73385924 | + | CFL | X | | Region not found | | | |
| X | 102642308-102661464 | + | CFL | X | 101153588-101170718 | CFL | | | 98.76 |
| X | 103734398-103744217 | + | CFL | X | | Region not found | | | 96.1 |
| X | 123894822-123902652 | + | pol, env | X | 122367428-122373579 | pol, env | | | 99.805 |
| X | 124135377-124143863 | + | gag, pro, env | X | 122608017-122616071 | gag, pro, env | Duplication of TM | | 86.841 |
| X | 124248563-124259123 | + | CFL | X | 122693788-122696469 | Solo | Homologous recombinatio n | | 99.238 |
| X | 138438214-138448602 | + | gag, pro, env | X | 137085311-137093829 | gag, pro, env | | | 99.191 |
| X | 139222079-139235419 | + | gag, pro, env | X | 137497621-137509492 | gag, pro, env | | Duplication and inversion (X: 137497621-137509492 , X: 139222079-139235419) | 93.591 |

| | | | | | | | | | |
|---|---|---|---|---|---|---|---|---|---|
| X | 173973872-173982128 | + | gag, pol, env | X | 171754474-171760827 | gag, pol, env | | | 98.541 |
| X | 48987788-48990506 | - | 5' Truncated | X | 48902687-48905404 | 5' Truncated | | | 99.963 |
| X | 100110589-100113302 | - | 5' Truncated | X | 98632445-98635158 | 5' Truncated | | | 99.816 |
| X | 108442887-108445593 | - | 5' Truncated | X | 107030410-107033116 | 5' Truncated | | | 100 |
| X | 116994594-116997312 | - | 5' Truncated | X | 115558538-115561256 | 5' Truncated | | | 100 |
| X | 117016876-117019589 | - | 3' Truncated | X | 115580712-115583420 | 3' Truncated | | | 98.822 |
| X | 138481572-138484274 | - | 3' Truncated | X | 137138488-137141191 | 3' Truncated | | | 99.593 |
| X | 149194002-149196677 | - | 5' Truncated | X | 147779246-147781918 | 5' Truncated | | | 99.568 |
| X | 154301915-154304627 | - | 3' Truncated | X | 152901855-152904567 | 3' Truncated | | | 100 |
| X | 156283681-156286399 | - | 5' Truncated | X | 154904080-154906798 | 5' Truncated | | | 99.89 |
| X | 156305596-156308311 | - | 3' Truncated | X | 154925991-154928706 | 3' Truncated | | | 99.963 |
| X | 74913305-74916018 | + | 5' Truncated | X | 73532419-73535132 | 5' Truncated | | | 99.926 |
| X | 75393858-75396561 | + | 5' Truncated | X | 74015476-74018179 | 5' Truncated | | | 100 |

| | | | | | | | | | |
|---|---|---|---|---|---|---|---|---|---|
| X | 81835813-81838523 | + | 5' Truncated | X | 80674106-80676818 | 5' Truncated | | | 97.722 |
| X | 88613139-88615854 | + | 5' Truncated | X | 87123551-87126266 | 5' Truncated | | | 100 |
| X | 98700867-98703593 | + | 3' Truncated | X | 97213729-97216455 | 3' Truncated | | | 99.936 |
| X | 120815899-120818610 | + | 5' Truncated | X | 119378570-119381281 | 5' Truncated | | | 99.926 |
| X | 151471419-151474120 | + | 5' Truncated | X | 150090000-150092701 | 5' Truncated | | | 99.963 |
| X | 156646436-156649142 | + | Solo | X | 155272152-155274103 | Two solo | Duplication of LTR | | 100 |

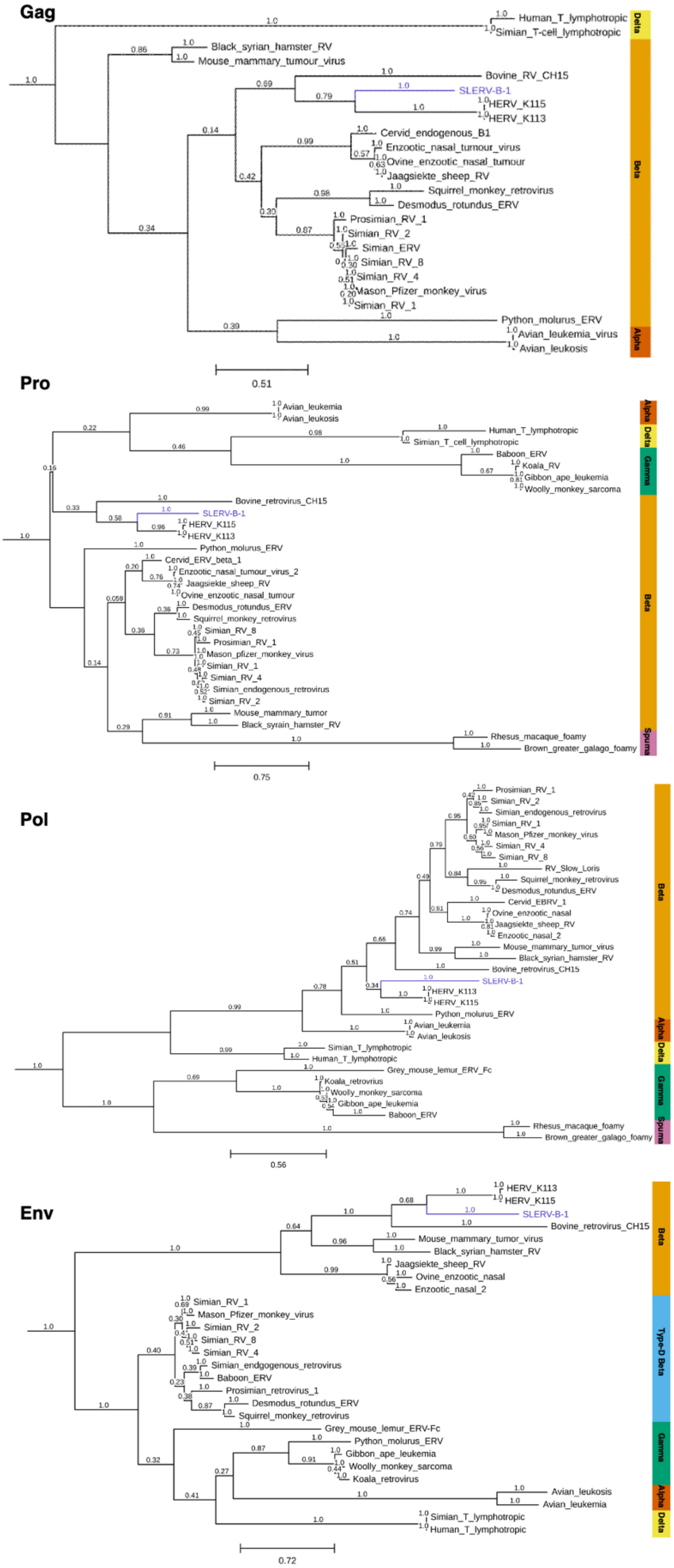


**Supplementary Figure 2: Phylogeny of SLERV-B-1 (in purple) inferred from Maximum**

**Likelihood (PhyML) analysis with bootstrap support shown. Scale bar represents substitutions per site. Trees have been rooted at midpoint**

**Supplementary table 4: Additional sequences used for phylogenetic reconstruction of *N. bengalensis* and *N. coucang* from Blair et al., (2023) and Pozzi et al., (2015).**

| Species | Genbank ID | Gene |
|---|---|---|
| *Nycticebus bengalensis* | KP410608 | Cytochrome B |
| *Nycticebus bengalensis* | KP410618 | Cytochrome B |
| *Nycticebus bengalensis* | KP410638 | Cytochrome B |
| *Nycticebus bengalensis* | KP410649 | Cytochrome B |
| *Nycticebus bengalensis* | KP410652 | Cytochrome B |
| *Nycticebus bengalensis* | KP410662 | Cytochrome B |
| *Nycticebus bengalensis* | KP410666 | Cytochrome B |
| *Nycticebus bengalensis* | OQ555560 | Cytochrome B |
| *Nycticebus bengalensis* | OQ555561 | Cytochrome B |
| *Nycticebus bengalensis* | OQ555565 | Cytochrome B |
| *Nycticebus coucang* | KP410591 | Cytochrome B |
| *Nycticebus coucang* | KP410592 | Cytochrome B |
| *Nycticebus coucang* | KP410655 | Cytochrome B |
| *Nycticebus coucang* | OQ555566 | Cytochrome B |
| *Nycticebus javanicus* | OQ555567 | Cytochrome B |
| *Nycticebus bengalensis* | NC_021958 | Cytochrome B, NADH dehydrogenase subunit 4, Cytochrome C oxidase subunit I |
| *Nycticebus coucang* | NC_002765 | Cytochrome B, NADH dehydrogenase subunit 4, Cytochrome C oxidase subunit I |
| *Nycticebus bengalensis* | OQ518050 | Cytochrome C oxidase subunit I |
| *Nycticebus bengalensis* | OQ518051 | Cytochrome C oxidase subunit I |
| *Nycticebus bengalensis* | OQ518052 | Cytochrome C oxidase subunit I |
| *Nycticebus bengalensis* | OQ518071 | Cytochrome C oxidase subunit I |
| *Nycticebus bengalensis* | OQ518072 | Cytochrome C oxidase subunit I |
| *Nycticebus bengalensis* | OQ518073 | Cytochrome C oxidase subunit I |
| *Nycticebus bengalensis* | OQ518074 | Cytochrome C oxidase subunit I |
| *Nycticebus bengalensis* | OQ518075 | Cytochrome C oxidase subunit I |
| *Nycticebus bengalensis* | OQ518076 | Cytochrome C oxidase subunit I |
| *Nycticebus bengalensis* | OQ518077 | Cytochrome C oxidase subunit I |

| | | |
|---|---|---|
| *Nycticebus bengalensis* | OQ518078 | Cytochrome C oxidase subunit I |
| *Nycticebus bengalensis* | OQ518079 | Cytochrome C oxidase subunit I |
| *Nycticebus bengalensis* | OQ518085 | Cytochrome C oxidase subunit I |
| *Nycticebus bengalensis* | OQ518086 | Cytochrome C oxidase subunit I |
| *Nycticebus bengalensis* | OQ518087 | Cytochrome C oxidase subunit I |
| *Nycticebus bengalensis* | OQ518088 | Cytochrome C oxidase subunit I |
| *Nycticebus coucang* | OQ518053 | Cytochrome C oxidase subunit I |
| *Nycticebus coucang* | OQ518054 | Cytochrome C oxidase subunit I |
| *Nycticebus coucang* | OQ518055 | Cytochrome C oxidase subunit I |
| *Nycticebus coucang* | OQ518055 | Cytochrome C oxidase subunit I |
| *Nycticebus coucang* | OQ518056 | Cytochrome C oxidase subunit I |
| *Nycticebus coucang* | OQ518056 | Cytochrome C oxidase subunit I |
| *Nycticebus coucang* | OQ518057 | Cytochrome C oxidase subunit I |
| *Nycticebus coucang* | OQ518057 | Cytochrome C oxidase subunit I |
| *Nycticebus coucang* | OQ518058 | Cytochrome C oxidase subunit I |
| *Nycticebus coucang* | OQ518058 | Cytochrome C oxidase subunit I |
| *Nycticebus coucang* | OQ518059 | Cytochrome C oxidase subunit I |
| *Nycticebus coucang* | OQ518060 | Cytochrome C oxidase subunit I |
| *Nycticebus coucang* | OQ518061 | Cytochrome C oxidase subunit I |
| *Nycticebus coucang* | OQ518062 | Cytochrome C oxidase subunit I |
| *Nycticebus coucang* | OQ518063 | Cytochrome C oxidase subunit I |
| *Nycticebus coucang* | OQ518064 | Cytochrome C oxidase subunit I |
| *Nycticebus coucang* | OQ518065 | Cytochrome C oxidase subunit I |
| *Nycticebus coucang* | OQ518066 | Cytochrome C oxidase subunit I |
| *Nycticebus coucang* | OQ518067 | Cytochrome C oxidase subunit I |
| *Nycticebus coucang* | OQ518068 | Cytochrome C oxidase subunit I |
| *Nycticebus coucang* | OQ518069 | Cytochrome C oxidase subunit I |
| *Nycticebus coucang* | OQ518070 | Cytochrome C oxidase subunit I |
| *Nycticebus javanicus* | OQ518082 | Cytochrome C oxidase subunit I |
| *Nycticebus bengalensis* | OQ555481 | NADH dehydrogenase subunit 4 |
| *Nycticebus bengalensis* | OQ555482 | NADH dehydrogenase subunit 4 |

| | | |
|---|---|---|
| *Nycticebus bengalensis* | OQ555483 | NADH dehydrogenase subunit 4 |
| *Nycticebus bengalensis* | OQ555484 | NADH dehydrogenase subunit 4 |
| *Nycticebus bengalensis* | OQ555492 | NADH dehydrogenase subunit 4 |
| *Nycticebus bengalensis* | OQ555496 | NADH dehydrogenase subunit 4 |
| *Nycticebus bengalensis* | OQ555497 | NADH dehydrogenase subunit 4 |
| *Nycticebus bengalensis* | OQ555519 | NADH dehydrogenase subunit 4 |
| *Nycticebus bengalensis* | OQ555520 | NADH dehydrogenase subunit 4 |
| *Nycticebus bengalensis* | OQ555532 | NADH dehydrogenase subunit 4 |
| *Nycticebus bengalensis* | OQ555539 | NADH dehydrogenase subunit 4 |
| *Nycticebus coucang* | OQ555475 | NADH dehydrogenase subunit 4 |
| *Nycticebus coucang* | OQ555477 | NADH dehydrogenase subunit 4 |
| *Nycticebus coucang* | OQ555478 | NADH dehydrogenase subunit 4 |
| *Nycticebus coucang* | OQ555485 | NADH dehydrogenase subunit 4 |
| *Nycticebus coucang* | OQ555486 | NADH dehydrogenase subunit 4 |
| *Nycticebus coucang* | OQ555488 | NADH dehydrogenase subunit 4 |
| *Nycticebus coucang* | OQ555489 | NADH dehydrogenase subunit 4 |
| *Nycticebus coucang* | OQ555490 | NADH dehydrogenase subunit 4 |
| *Nycticebus coucang* | OQ555491 | NADH dehydrogenase subunit 4 |
| *Nycticebus javanicus* | OQ555474 | NADH dehydrogenase subunit 4 |
